# From Individual to Shared Tempo: How Spontaneous Tempo Preferences Impact Joint Performance

**DOI:** 10.1101/2025.09.14.676070

**Authors:** Leah Snapiri, Yael Kaplan, Ayelet N. Landau

**Affiliations:** Department of Psychology, The Hebrew University of Jerusalem; Department of Cognitive Science, The Hebrew University of Jerusalem; Department of Experimental Psychology, University College London

## Abstract

Endogenous rhythmic structure is a key characteristic of our cognitive system, organizing both sensory processes and motor behavior. In the motor domain, recent studies demonstrated that individuals vary systematically in the spontaneous tempo of self-paced movement, exhibiting consistent tempo preferences across time, tasks, and cognitive demands. In our daily experience, we often need to perform with others, adjusting our motor signatures to support interpersonal coordination. The purpose of this study is to examine how individuals with distinct spontaneous tempi reconcile their personal tempo preferences with the need to synchronize during joint performance. To this end, we measured individuals’ spontaneous tempo across different days using three motor behaviors: tapping, walking, and bouncing. This design allowed us to examine both temporal consistency and cross-task consistency. Participants were then asked to perform these tasks together under two experimental conditions: (1) side-by-side, without explicit instructions to synchronize, and (2) with explicit instructions to synchronize their movements. Our findings replicate and extend recent work, demonstrating that individuals show stable rhythmic preferences across time, tasks, and interactive contexts. When performing together, two complementary patterns emerged. First, dyads converged on a common tempo around 2 Hz, reflecting a preferred tempo for joint rhythmic performance. Second, individuals retained a bias toward their own spontaneous tempo during joint performance, with faster or slower individuals performing correspondingly fast or slow within the range of dyadic tempo. Finally, individuals with more similar spontaneous tempi achieved greater synchrony and temporal stability when performing together. Thus, while individuals can flexibly adjust their spontaneous tempo to support joint action, personal tempo preferences remain a robust source of variability in how people perform and interact with others.

## Introduction

Internally generated rhythms are a hallmark of biological systems (Bunning, 1956), appearing across sensory (Baumgarten et al., 2015; VanRullen, 2016; Zoefel & VanRullen, 2017), cognitive (Fiebelkorn & Kastner, 2019; Herbst & Landau, 2016; Re et al., 2019, 2025) and motor domains (Amit et al., 2017; Benedetto et al., 2020; Berg, 2002; Melloni et al., 2009). In the motor domain, endogenous rhythmic properties are often assessed through self-paced rhythmic motion, such as during a simple tapping or walking task (Fraisse, 1982; McAuley et al., 2006; Rimoldi, 1951). Although pioneering studies have empathized the homogeneity of spontaneous motor rhythms (Fraisse, 1982; McAuley, 2010; Moelants, 2002), and individual differences were tested in the context of life-span development (McAuley et al., 2006), musical skill (Scheurich et al., 2018) and clinical populations (Kliger Amrani & Zion Golumbic, 2020). More recent work suggests that substantial and systematic variability can be observed among healthy adults (Desbernats et al., 2023; Engler et al., 2024). For example, Engler and colleagues (2024) explored the expression of endogenous rhythms through spontaneous, self-paced movement and found that spontaneous tempi are highly consistent across measurement days and generalize across movement types, including tapping, walking, and clapping. Furthermore, this consistency was observed across tasks with varying cognitive demands, from simple tapping to complex musical production. Together, these findings suggest that spontaneous motor tempo reflects a stable motor trait, that captures individual differences in endogenous rhythmic properties.

This raises a fundamental question: what happens to these stable endogenous rhythms when people perform together? In everyday life, coordination often involves adapting to the rhythmic structure generated by others, whether during a conversation, walking side-by-side, or making music. This process requires balancing individual tempo preferences with the emerging demands of shared timing. How do individuals with distinct spontaneous tempi coordinate effectively during joint performance? In a pivotal study on the movement of fish fins, von Holst (1973) observed that while each fin tends to oscillate at a specific frequency when moving individually, when two fins oscillate stimulatory they exhibit a mutual “magnet effect”. Namely, each fin pulls the other toward its own preferred frequency, resulting in convergence at an intermediate frequency. De Rugy and colleagues (2006) tested whether similar principles apply when two individuals coordinate by oscillating handheld pendulums. To this end, they experimentally manipulated each participant’s preferred frequency by adjusting the length and weight of the pendulums. They found that the joint tempo typically converged to a value between the two preferred frequencies. Furthermore, they showed that as asymmetries between partners increased, due to differences in the physical properties of the pendulums, synchrony decreased. These results show that successful coordination emerges through dynamic negotiation and is modulated by each participant’s intrinsic movement tempo.

Extending previous research on coordination dynamics, the present study shifts the focus from experimentally manipulated preferred movement frequencies to intrinsic individual differences. By measuring each participant’s spontaneous motor tempo, we investigate how the natural variability between individuals shapes the quality of interpersonal coordination. To this end, participants completed two experimental sessions involving three simple motor tasks: tapping, walking, and bouncing. In the first session, participants performed each task individually to assess their spontaneous tempo, defined as the mean inter-onset interval (IOI) between consecutive events. In the second session, the same tasks were repeated under three conditions: (1) individually, to assess consistency in spontaneous tempi over time (2) jointly, without explicit instructions to synchronize, and (3) jointly, with explicit instructions to synchronize. We ask how individual spontaneous tempi are modulated during deliberate coordination. Additionally, we examine whether tempo similarity between partners, measured as the absolute difference between their mean IOIs, predicts the quality of joint performance. Through this work, we aim to further understand how endogenous motor rhythms shape, and are shaped by interpersonal interaction, and shed light on the interplay between individuality and togetherness during simple joint tasks.

## Methods

### Participants

The experiment consisted of two experimental sessions conducted on different days. Seventy participants completed the first experimental session (77.14 % females, 90% right-handed, mean age = 22.33 [SD = 3.12]). Fifty-two participants (76.92% females, 92.31% right-handed, mean age = 22.81[SD = 3.18]) completed both experimental sessions. Participants were matched to perform together in the dyadic session based on gender and native language (65.38% Hebrew, 19.23% Arabic, 7.69% English, 7.69% other). We obtained digital informed consent from all individuals before the experimental sessions. All experimental procedures were approved by the local Ethics Committee of the Hebrew University of Jerusalem, Israel.

### Spontaneous Motor Tempo Across Time and Tasks

To characterize individuals’ spontaneous motor tempo, we asked participants to perform three simple motor tasks: tapping, walking, and bouncing. We measured the spontaneous tempo with which individuals perform these tasks across two experimental sessions conducted on different days. We instructed participants to perform these tasks at a comfortable and regular tempo.

### Tapping task

Participants tapped with the index finger of their dominant hand on a smartphone screen for one minute. The timing of each tap was recorded using a touch-sensitive app developed in the lab. The smartphone was placed on a table in front of the participant at a comfortable distance.

### Walking task

Participants walked for three minutes along a long, circular corridor adjacent to the experimental area. The timing of each step was recorded using a motion-sensitive app based on the smartphone’s accelerometer and GPS sensors. The phone was attached to the participant’s upper arm using an armband. Before the recording began, participants completed a short practice walk to calibrate the sensor to their movement characteristics.

### Bouncing task

Participants bounced on a 75 cm Pilates ball for three minutes at a regular and comfortable pace. Bounce times were recorded using a motion-sensitive app based on the accelerometer and gyroscope sensors of the smartphone. Similarly to the walking task, the phone was attached to the participant’s upper arm using an armband and a brief practice was conducted prior to recording to ensure proper sensor calibration. All behaviors were recorded using lab-developed Android applications installed on Xiaomi Redmi Note 5 smartphones.

### Spontaneous Tempi in Joint Action

To examine how individual tempo preferences impact joint rhythmic action we paired participants into dyads and asked them to perform the three motor tasks together under two distinct conditions.

### Side-by-side condition

Participants were instructed to perform the task simultaneously at a tempo that felt natural and comfortable, without any requirement to synchronize. In the tapping task, participants sat across from each other at the same table, each performing the task on a separate lab-provided smartphone. They tapped for one minute, with start and end times cued by the experimenter. In the walking task, participants walked side by side along a corridor for three minutes. In the bouncing task, participants bounced on Pilates balls for three minutes while facing each other at a comfortable distance in the experimental area. Across all tasks, participants were asked not to speak during performance.

### Deliberate coordination condition

The setup was identical to the side-by-side condition. However, participants were instructed to actively coordinate their movements and synchronize as accurately as possible.

### Procedure

Participants provided informed consent and completed a demographic questionnaire prior to the first experimental session. Upon arrival, they completed an individual assessment of spontaneous motor tempo. Each participant performed the motor tasks in a fixed order: (1) tapping, (2) walking, and (3) bouncing. This initial assessment lasted 15 minutes. Participants were invited for a second session following a week to one month after the first. This session began with a repeated individual assessment to examine the consistency of spontaneous motor tempi over time. Then, participants were introduced to an assigned partner (another returning participant) and performed all three motor tasks again, this time as a dyad. Participants completed the side-by-side condition first, then the synchronized condition. The second session lasted 60 minutes. For an overview see Figure 1.

**Figure 1.**
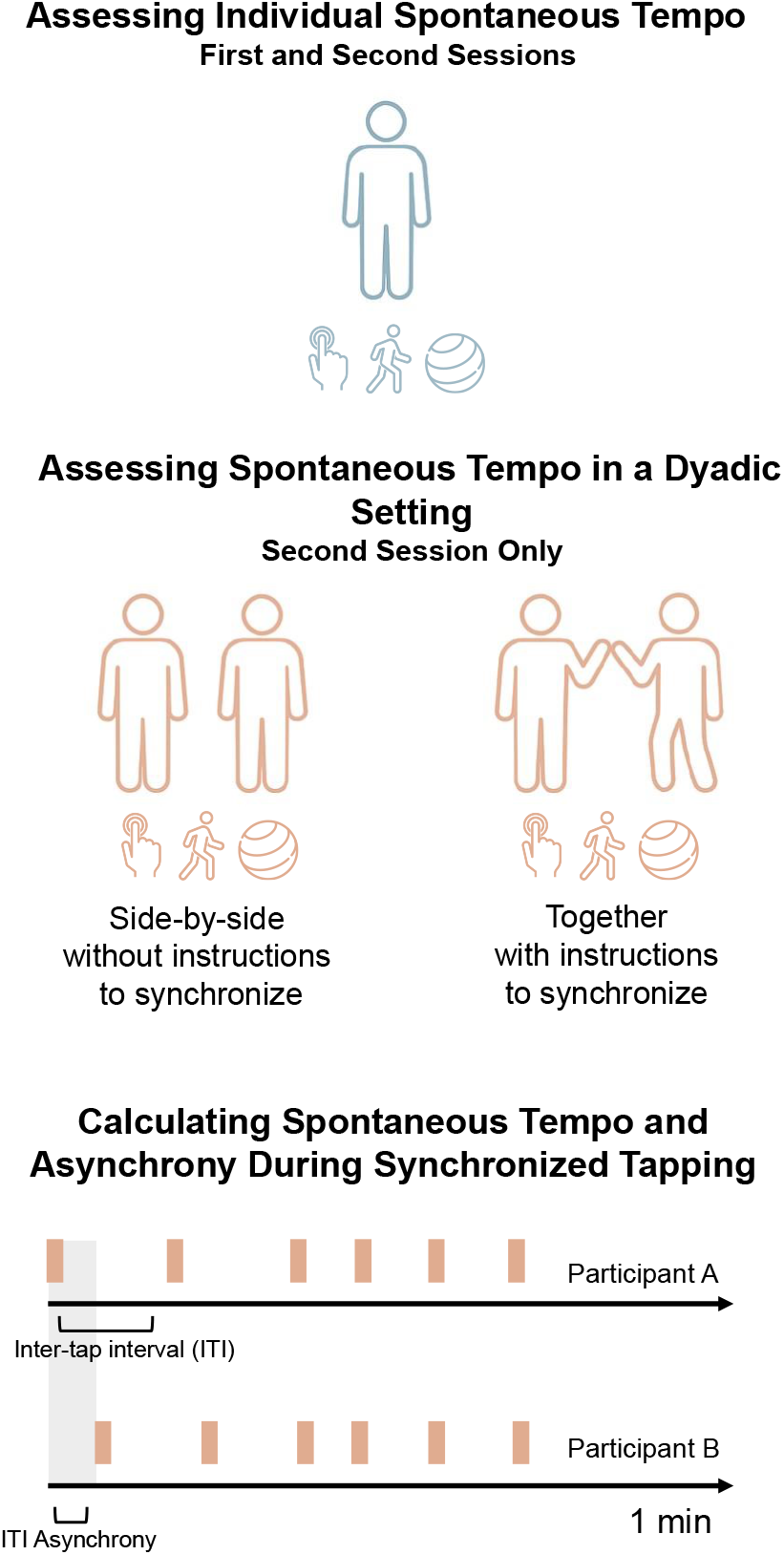
Overview of experimental design and key parameters. Participants completed two experimental sessions. In the first session, they performed three motor tasks: tapping (1 min), walking (3 min), and bouncing (3 min), individually. In the second session, participants were paired into dyads and repeated all three tasks under three conditions: individually, side by side with a partner, and together with explicit instructions to synchronize. For each participant and task, we calculated spontaneous tempo based on the mean inter-onset interval (IOI) between successive events (taps, steps, or bounces), yielding 12 measurements of spontaneous tempo per participant (3 tasks × 4 repetitions). The bottom panel illustrates how we computed inter-tap-interval (ITI) asynchrony between two participants during synchronized tapping. Taps are indicated by red rectangles, and asynchronies were calculated based on the absolute difference between the participants’ ITIs throughout the duration of the task.

## Data Analysis

### Assessing Spontaneous Motor Tempo

For each participant and measurement, we calculated the inter-onset intervals (IOIs) between recorded events (i.e., between taps, steps, and bounces) and pre-processed the time course of performance using the following steps: (1) IOIs longer than 3 seconds were removed, as these were indicative of breaks in performance; (2) IOIs that were more than 1.5 interquartile ranges above the upper quartile or below the lower quartile were excluded. On average, these procedures resulted in the removal of 3.76% of taps (SD = 2.49), 3.96% of steps (SD = 4.08), and 3.26% of bounces (SD = 2.48), per measurement.

Then, for each measurement, we calculated two performance measures: (1) **Spontaneous tempo**, defined as the mean IOI between recorded events; and (2) **Coefficient of variation (CV)**, a measure of performance variability that accounts for individual differences in spontaneous tempo, by dividing the standard deviation of inter-onset intervals by the mean. Participants with extreme CV or spontaneous tempo values >3.5 SD from the mean were excluded, removing 3% of participants per measurement on average (SD = 1.35).

### Assessing Interpersonal Coordination During Joint Action

To evaluate the quality of interpersonal coordination, we focused on the tapping task performed under the condition of deliberate coordination. For each dyad, we calculated IOI Asynchrony by taking the difference between partners’ inter-onset intervals (IOIs) for each corresponding tap. The absolute values of these differences were then averaged across the entire performance, yielding an overall index of temporal misalignment (for an illustration see Figure 1). In addition to assessing between-partner coordination, we examined within-participant temporal stability. For each individual, we computed the standard deviation of their IOIs during the synchronized tapping task. These variability values were then averaged within dyads, providing a measure of how consistently each dyad maintained a stable tempo throughout performance. In all coordination analyses, we used raw timing data without excluding events, in order to capture missed taps and breaks in synchrony. Participants were also not excluded based on high variability or extreme tempi, as our focus was on the quality of joint performance rather than specific tempo characteristics. This approach allowed us to include a wider range of natural coordination behavior, including atypical or unstable performances. Additionally, due to logging failures, we did not record spontaneous tempo during the first session for two participants. For these individuals, we used their second session tapping data to estimate spontaneous tempo distance from their partner.

## Results

### Variability in Spontaneous Motor Tempi Across Tasks

As shown in Figure 2, participants exhibited a wide range of spontaneous tempi when tapping, with mean inter-onset intervals (IOIs) ranging from 0.170 to 1.370 seconds (M = 0.567 sec, SD = 0.257 sec). In contrast, spontaneous walking and bouncing tempi varied less, with walking IOIs ranging from 0.421 to 0.761 seconds (M = 0.571 sec, SD = 0.060 sec) and bouncing IOIs from 0.637 to 0.860 seconds (M = 0.710 sec, SD = 0.040 sec). Correspondingly, between-subject variability was substantially higher than within-subject variability for tapping (between-subject CV = 0.45; within-subject mean CV = 0.07), but not for walking (CV = 0.10 vs. 0.07), or bouncing (CV = 0.06 vs. 0.03).

**Figure 2.**
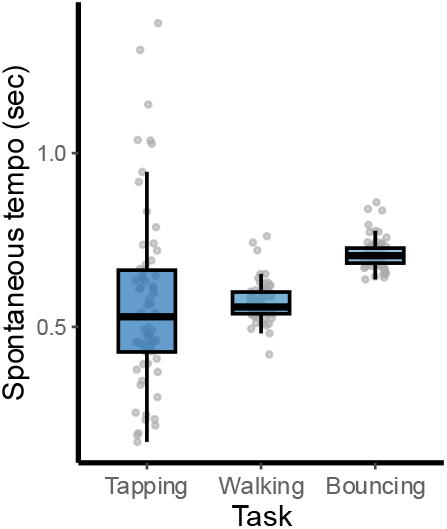
Variability in spontaneous tempi is task dependent. Distributions of spontaneous motor tempi across the three motor tasks as measured during the first solo assessment. Individuals showed substantially greater variability in spontaneous tapping tempi, compared to walking and bouncing.

### Spontaneous Tempo is a Stable Individual Trait

We examined the consistency of spontaneous tempo preferences across time, tasks, and interactive settings using Spearman correlations. We used spearman correlation to constrain the impact of extreme observations on correlation values and address deviations from homoscedasticity. Statistical tests were one-sided, with an alpha level of 0.05. Reported p- values were adjusted for multiple comparisons using the False Discovery Rate (FDR) correction method (Benjamini et al., 2001).

Individuals showed stable spontaneous tempi over time, as reflected in highly significant test–retest correlations: tapping, r_s_ (45) = .82; walking, r_s_ (44) = .66; and bouncing, r_s_ (44) = .50 (all *p* < .001; see Figure 3a). Furthermore, Spontaneous tapping tempo significantly predicted walking tempo, in both experimental sessions (**session 1:** r_s_ (61) = .25, *p* = .03, **session 2**: r_s_ (45) = .34, *p* = .01, Figure 3b). However, bouncing tempo was not correlated with either tapping or walking tempi. Finally, individuals’ spontaneous tempo measured in the first solo session reliably predicted their tempo during the side-by-side condition in the second session across all three motor behaviors (**tapping**: r_s_ (48) = .52, *p* < .001, **walking:** r_s_ (46) = .59, *p* < .001, and **bouncing**: r_s_ (47) = .49, *p* < .001, Figure 3c). Therefore, individuals exhibit stable tempo preferences that persist across time and social contexts. Among the three movement types, tapping exhibited the greatest inter-individual variability, the highest test-retest reliability, and significantly predicted both walking tempo and spontaneous tempo during side-by-side performance. Accordingly, we focused our subsequent analyses on the tapping task, to examine how individual tempo preferences influence interpersonal coordination across different interactive settings.

**Figure 3.**
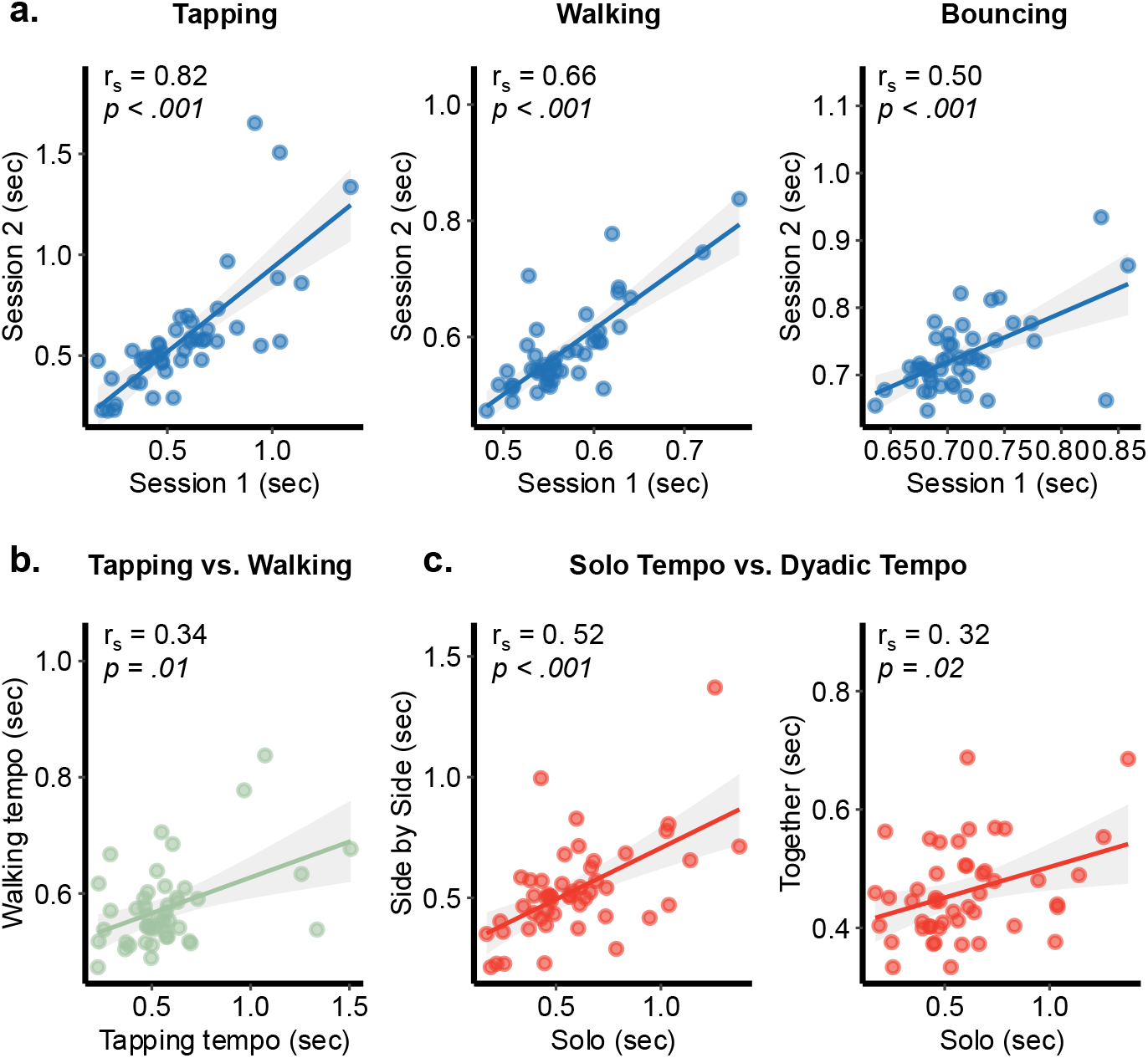
Spontaneous motor tempi reflect a consistent individual trait. **(a)** Individuals showed highly consistent spontaneous motor tempi across two measurement days, for all motor behaviors: tapping, walking, and bouncing. The top three panels plot each participant’s mean inter-onset interval (IOI, in seconds) from the first session (x-axis) against the second session (y-axis). Each point represents a single participant. Spearman correlation coefficients and FDR-corrected p-values are shown for each task, indicating strong test–retest reliability. **(b) Individual spontaneous tempo preferences were consistent across different motor behaviors**. The plot shows the mean inter-onset intervals (IOIs) for tapping (x-axis) and walking (y-axis) from the second solo session. Individuals’ tapping tempo significantly predicted their walking tempo. Namely fast tappers tended to walk relatively fast and slow tappers tended to walk relatively slow, within the restricted range of walking tempi. This relationship was observed in both experimental sessions. **(c) Individual tapping tempo during the first solo session (x-axis) predicted tapping tempo during both dyadic conditions**: left panel shows the side-by-side condition, and right panel shows the deliberate synchronization condition. This relationship between solo assessment and dyadic assessments was observed also for walking and bouncing.

### A Common Tempo of Joint Performance Across Dyads

We next asked how coordinating with a partner affects individuals’ spontaneous motor tempo. As participants tended to maintain their spontaneous tempo during the side-by-side condition we focus here on dyadic performance during deliberate synchronization. We first conducted a paired-samples t-test to evaluate changes in spontaneous tapping tempi between the solo tapping task during the first session and the joint synchronized tapping during the second session. On average, individuals tapped at a faster tempo during joint performance compared to their spontaneous tapping tempo during the solo assessment (mean difference = −0.14 sec, 95% CI [−0.21, −0.06], *t* (46) = −3.61, *p* < .001; Cohen’s *d* = –0.53, 95% CI −0.84, −0.22]). However, this shift was not consistent across participants and varied systematically with their individual spontaneous tempo (see Figure 4b). Specifically, individuals with slower spontaneous tempi during the solo session (i.e., longer ITIs, above the median) tended to speed up during joint tapping, whereas individuals with faster spontaneous tempi (below the median) either slowed down or showed minimal change. Together, these opposing adjustments led to a pronounced convergence toward a shared tempo during dyadic performance, centered around an inter-tap interval of 0.460 seconds (SD = 0.08). Notably, even dyads in which both participants had similar spontaneous tempi, whether fast or slow, converged toward this common dyadic tempo during joint performance (Figure 4a). This convergence point (~2 Hz) aligns closely with the median spontaneous tempo in the population (Moelants, 2002) and has been identified as a “sweet spot” for sensorimotor synchronization (McAuley et al., 2006; Repp & Su, 2013). These findings suggest that individuals implicitly adjust their spontaneous tempo preferences toward a commonly optimal tempo to facilitate interpersonal synchrony.

**Figure 4.**
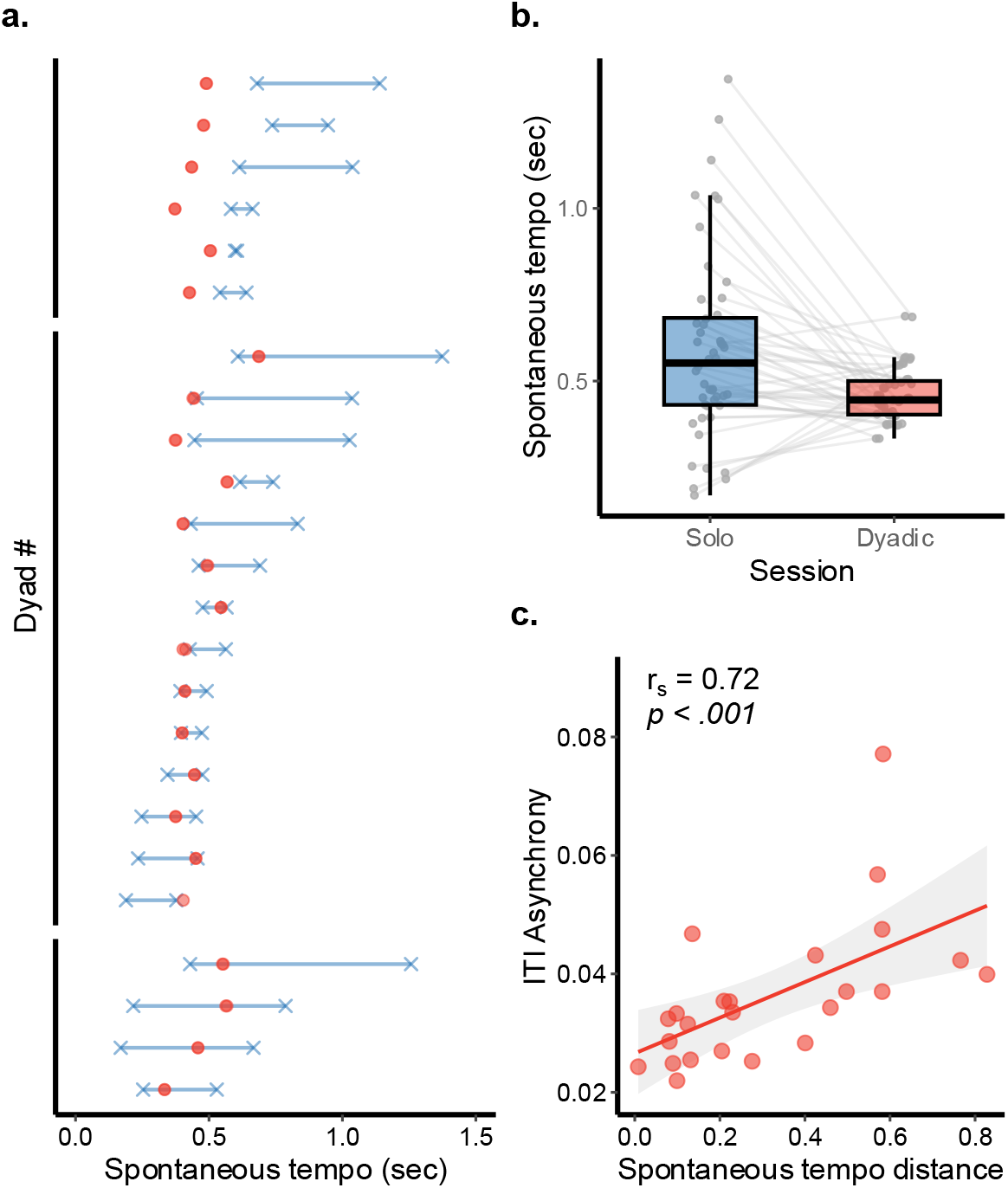
**(a) Spontaneous motor tempi dynamically shift to support interpersonal coordination**. Blue crosses indicate each participant’s spontaneous tempo during the first solo session (mean inter-tap interval (ITI) measured in seconds). Red circles indicate the tempo produced during deliberate synchronization with a partner. The y-axis represents individual dyads, sorted within each panel by their average spontaneous tempo, as measured during the first solo session. Dyads are grouped into three panels to show that participants reached a common tempo through varied coordination strategies and from diverse set points. Bottom panel: participants met halfway by shifting toward each other’s tempo. Middle panel: one participant adjusted while the other maintained their spontaneous tempo. Top panel: Both participants sped up with respect to their individual spontaneous tempo to reach the common dyadic tempo. Note that in most dyads, the two red circles overlap, reflecting successful synchronization on average. In one dyad (fifth from the bottom), only one red circle is shown due to a missing measurement. **(b) Joint performance is characterized by reduced variability in spontaneous tempi**. Participants showed substantial individual differences in spontaneous tapping tempo during the solo session, reflected in the wide distribution of IOIs (blue boxplot). During deliberate synchronization, however, they converged toward a shared tempo, as shown by the narrower distribution centered around 0.460 seconds (red boxplot). **(c) Individuals with more similar spontaneous tempi synchronized more effectively**. The x-axis shows spontaneous tempo distance within each dyad, calculated as the absolute difference between the spontaneous tempi of the two participants, as measured during the first solo session. The y-axis shows mean ITI asynchrony during joint tapping, calculated as the average of the absolute differences between the inter-tap intervals of the two participants during deliberate synchronization. As spontaneous tempo distance increased, so did asynchrony. Spearman correlation coefficient and corresponding p-value are shown in the upper left corner of the figure.

### Individual Tempo Preferences Impact Dyadic Tempo and Synchronization Quality

To assess how individual spontaneous tempo preferences influence joint performance we first tested the relationship between each participant’s spontaneous tempo, as measured during the solo session, and their tempo during the dyadic session. Although participants converged toward a common tempo when tapping together, their dyadic tempo was systematically linked to their solo tempo preferences. Specifically, individuals with faster spontaneous tempi tended to tap faster than others during joint tapping, while those with slower tempi remained relatively slower (r_s_ (46) = .32, *p* = .02, Figure 3c).

Next, we examined whether tempo similarity between partners affected the quality of their synchronization. For each dyad, we calculated **spontaneous tempo distance** as the absolute difference between their mean inter-tap intervals (ITIs) from the solo session. We then calculated the mean ITI asynchrony between the two participants during joint tapping (see Methods). We found that spontaneous tempo distance is positively linked to ITI synchrony (r_s_ (22) = .72, *p* < .001, Figure 4c). Namely, individuals with similar spontaneous tempi had lower ITI asynchrony values. Finally, we tested whether spontaneous tempo distance predicted variability in tapping tempo during joint performance. We found the variability in joint tapping increased with greater spontaneous tempo distance between participants (r_s_ (22) = .65, *p* = .001). Therefore, individuals with similar spontnapus tempi were better able to maintain a stable beat during the task. Taken together, these findings suggest that even as individuals adjust their tempo to achieve optimal synchronization, their spontaneous tempo preferences continue to influence both the tempo and quality of joint performance.

## Discussion

In the current study, we examined how individual differences in endogenous rhythms influence dyadic coordination during joint rhythmic action. To assess individuals’ endogenous rhythms, we used three rhythmic motor tasks: tapping, walking, and bouncing. Participants performed these tasks across different measurement days and across different interactive conditions: individually, side by side, and while actively synchronizing with a partner. We found that individuals were highly consistent in their spontaneous motor tempo across time and interactive conditions. We also observed significant cross-task correlations: individuals with faster spontaneous tapping tempi also tended to walk faster, and vice versa. In addition to this intra-individual consistency, we found that the extent of variability between individuals varied significantly across tasks. Tapping revealed substantial variation in spontaneous tempi across individuals, whereas walking and bouncing were more homogenous. This homogeneity likely reflects stronger biomechanical and physical constraints associated with walking and bouncing. In particular, walking tempo is influenced by factors like limb length and weight (Goodman et al., 2000; Nessler & Gilliland, 2009), while bouncing is additionally constrained by the physical properties of the Pilates ball.

Taken together, these findings suggest that the spontaneous tempo of movement reflects both domain-general endogenous rhythmic properties and task-specific constraints. While biomechanical and contextual factors may limit the range of tempi that can be expressed in a given task (Engler et al., 2024), the underlying rhythmic preferences remain stable across time, movement types and settings. This supports the view of spontaneous tempo as a consistent motor trait (Desbernats et al., 2023; Engler et al., 2024; Fraisse, 1982; Rimoldi, 1951). We next focused on spontaneous tapping, to examine how individual tempo preferences are modulated during interpersonal coordination.

Previous studies have shown that when individuals engage in joint rhythmic action, such as swinging hand-held pendulums, they tend to converge on a frequency that lies between the resonance frequencies of their respective pendulums (de Rugy et al., 2006; Schmidt & Turvey, 1989). This has been interpreted within a dynamical systems framework, where coupled oscillators with differing intrinsic frequencies naturally synchronize through mutual adaptation, often meeting halfway (Turvey et al., 1986; Von Holst, 1973). Our study revealed a contrasting pattern. When participants tapped together, they did not converge on an intermediate tempo between their individual spontaneous tempi. Instead, they collectively adopted a common tempo around 2 Hz (IOI = 0.500 sec), even when this tempo was substantially faster or slower than their individual preferences. These divergent results may reflect differences in task mechanics. In pendulum swinging, the physical properties of the pendulums may naturally favor synchronization at an intermediate frequency (Turvey et al., 1986). By contrast, joint tapping may be optimized around a specific, functionally advantageous tempo. Extensive research identifies 2 Hz as a “sweet spot” for sensorimotor synchronization (McAuley et al., 2006; Repp, 2006; Repp & Su, 2013). Moreover, this tempo corresponds to peak resonance in motor areas during action observation (Avanzino et al., 2015), and closely approximates the median spontaneous tempo observed in the general population (Hammerschmidt et al., 2021; Moelants, 2002). Thus, convergence at this tempo may reflect an implicit bias toward a rhythm that is both broadly familiar and functionally optimal for interpersonal coordination.

A similar pattern has been observed in a study on continuous unstructured motion during joint improvisation. While individuals display distinct motion characteristics when leading an interaction, they converge on shared motion characteristics during highly synchronized joint movement (Hart et al., 2014). Specifically, during moments of enhanced synchrony, motion trajectories become smooth and symmetric, often resembling half-period sine waves. This suggests a preference for motion patterns that are easy to predict and extrapolate. Therefore, when coordinating, individuals adopt movement patterns that enhance mutual predictability (Sacheli et al., 2013; Vesper et al., 2010, 2011). In our study, participants had no prior familiarity with each other’s movement characteristics (apart from the side-by-side condition), which may have led them to adopt highly predictable and broadly familiar tempo to facilitate synchronization. Future research could explore whether prior experience with a partner’s movement style, as in the case of close friends or frequent co-performers, allows for more flexible or idiosyncratic synchronization patterns.

In addition to converging on a shared tempo during joint performance, individuals showed a consistent bias toward their own spontaneous tempo. Those with faster or slower spontaneous tempi tended to tap at correspondingly faster or slower rates, even within the narrow range of dyadic tapping tempi. This finding extends previous research on solo synchronization tasks involving fixed external pacing cues (McAuley et al., 2006; Roman et al., 2023; Scheurich et al., 2018; Zamm et al., 2018). In tasks such as finger tapping or playing simple melodies with an external metronome, performance drifts back toward an individual’s spontaneous tempo when the pacing rhythm deviates substantially from it. Our results demonstrate that a similar pull toward spontaneous tempo occurs during interpersonal coordination, where the external rhythm is not imposed by a metronome but emerges through coordination with another person. This is consistent with entrainment-based models, which suggest that while individuals can adapt to a wide range of external tempi, their spontaneous tempo acts as an attractor state, continuously pulling behavior toward individuals’ tempo preferences (Jones & Boltz, 1989; Large & Jones, 1999; McAuley et al., 2006; Roman et al., 2023).

Finally, this study extends prior research on the role of spontaneous tempo similarity in shaping the quality of joint rhythmic performance. Previous work has shown that individuals with similar preferred movement frequencies synchronize more effectively, whether in studies that mechanically manipulated preferred movement frequencies (e.g., rocking chairs or pendulums, De Rugy et al., 2006; Richardson et al., 2007), or during joint musical performance tasks (Tranchant et al., 2022; Zamm et al., 2015, 2016). Consistent with these findings, we show that dyads with more similar spontaneous tapping tempi achieve greater synchrony and rhythmic stability. While dynamical systems framework explain this in terms of mutual adaptation between coupled oscillators, where similar intrinsic tempi reduce the effort required to synchronize (de Rugy et al., 2006; Heggli et al., 2019, 2021; Marsh et al., 2009; Roman et al., 2023), these results can also be understood through current models of rhythmic perception and performance. de Lafuente and colleagues (2024) trained monkeys to follow and internally maintain external visual rhythms at different tempi, revealing tempo-specific neural activity across a widespread network, including visual, parietal, premotor, prefrontal, and hippocampal areas. Both firing rates and broadband local field potentials oscillated at the rhythm’s tempo, both during external pacing and internal maintenance.

Relatedly, studies in action observation and motor imagery suggest that individuals understand and predict others’ actions in relation to their own motor repertoire (Calvo-Merino et al., 2006; Rizzolatti et al., 1996), including individual temporal characteristics. For example, during action observation individuals show maximal activation in motor areas when a rhythmic motion is performed at a tempo that is close to individuals’ spontaneous motor tempo (Avanzino et al., 2015). Taken together, these findings suggest that individuals’ internal neural dynamics, including endogenous rhythmic properties, can shape and constrain how individuals perform together during joint rhythmic action.

In summary, individuals maintained consistent spontaneous tempi across time, tasks, and interactive settings, reflecting a stable motor trait. When performing together, they flexibly adapted their spontaneous tempo to support optimal joint coordination. However, their underlying tempo preferences continued to influence performance in two main ways. First, participants tended to tap relatively faster or slower, according to their own spontaneous tempo, even when synchronizing with a partner. Second, dyads with more similar spontaneous tempi achieved greater synchrony and rhythmic stability. Therefore, spontaneous tempo represents a fundamental individual characteristic that impacts the quality and dynamics of joint rhythmic coordination. Future research could further examine whether individual differences in spontaneous tempo are similarly predictive of coordination in more complex, unstructured settings that resemble more closely everyday social interactions. Moreover, given the well-established links between interpersonal synchrony, social bonding, and positive affect (Hove & Risen, 2009; Marsh et al., 2009; Mogan et al., 2017; Ravreby et al., 2022), it will be interesting to explore how tempo similarity influences not only performance, but also the subjective experience of moving together.

